# Magnetothermal control of temperature-sensitive repressors in superparamagnetic iron nanoparticle-coated *Bacillus subtilis*

**DOI:** 10.1101/2022.06.18.496685

**Authors:** Emily M. Greeson, Cody S. Madsen, Ashley V. Makela, Christopher H. Contag

## Abstract

Superparamagnetic iron oxide nanoparticles (SPIONs) are used as contrast agents in magnetic resonance imaging (MRI) and magnetic particle imaging (MPI) and resulting images can be used to guide magnetothermal heating. Alternating magnetic fields (AMF) cause local temperature increases in regions with SPIONs, and we investigated the ability of magnetic hyperthermia to regulate temperature-sensitive repressors (TSRs) of bacterial transcription. The TSR, TlpA39, was derived from a Gram-negative bacterium, and used here for thermal control of reporter gene expression in Gram-positive, *Bacillus subtilis. In vitro* heating of *B. subtilis* with TlpA39 controlling bacterial luciferase expression, resulted in a 14.6-fold (12-hour; h) and 1.8-fold (1-h) increase in reporter transcripts with a 9.0-fold (12-h) and 11.1-fold (1-h) increase in bioluminescence. To develop magnetothermal control, *B. subtilis* cells were coated with three SPION variations. Electron microscopy coupled with energy dispersive X-ray spectroscopy revealed an external association with, and retention of, SPIONs on *B. subtilis*. Furthermore, using long duration AMF we demonstrated magnetothermal induction of the TSRs in SPION-coated *B. subtilis* with a maximum of 4.6-fold increases in bioluminescence. After intramuscular injections of SPION-coated *B. subtilis,* histology revealed that SPIONs remained in the same locations as the bacteria. For *in vivo* studies, 1-h of AMF is the maximum exposure due to anesthesia constraints. Both *in vitro* and *in vivo*, there was no change in bioluminescence after 1-h of AMF treatment. Pairing TSRs with magnetothermal energy using SPIONs for localized heating with AMF can lead to transcriptional control that expands options for targeted bacteriotherapies.

---

Magnetic nanoparticles have broad applications in biomedicine including imaging, drug delivery, theranostics and therapeutic hyperthermia.^1–3^ Nanoparticles have also been used to study and treat bacterial infections through the coating of bacterial membranes for imaging and as anti-microbial agents.^4–10^ Superparamagnetic iron oxide nanoparticles (SPIONs) are useful imaging contrast agents for magnetic resonance imaging (MRI) and more recently in magnetic particle imaging (MPI).^11–16^ MPI detects SPIONs directly, providing a readout of both iron content and location with high specificity and sensitivity.^11, 17–19^ Further, MPI can guide the application of electromagnetic energy generated by alternating magnetic fields (AMF) to cause local temperature increase known as magnetic hyperthermia^20–22^ to precisely heat the iron-containing area.^23^

*Bacillus subtilis* is a model Gram-positive organism^24^ with numerous synthetic biology strategies for manipulating gene expression,^25, 26^ global metabolic networks^27^ and the entire genome^28–30^ making it well-suited for engineering systems for spatial and temporal regulation.^27^ *B. subtilis* is a generally recognized as safe organism that is used for industrial protein production and is highly resistant to environmental stressors such as heat with a heat shock response at 48°C.^31, 32^ High heat resistance and well characterized protein production pathways may make *B. subtilis* an ideal chassis organism for thermal energy-controlled protein production that could act as therapeutics.^33–35^ *B. subtilis* also has multiple characterized inducible systems including several sugar-regulated inducible systems.^36^ *B. subtilis* has been well studied for a variety of *in vitro* industry applications in areas such as pharmaceutical/nutraceutical production, recombinant protein production and secretion, and production of functional peptides and oligopeptides.^37–40^ However, these inducible systems have limited control for both *in vitro* and *in vivo* applications due to potential host toxicity, cost and carbon-source dependence.^41^

Temperature-sensitive repressors (TSRs) are a class of repressors that bind an operator-promoter region with temperature dependence and show promise for *in vivo* control with local heating for localized delivery.^42^ With the addition of thermal energy to the system, a structural change occurs that releases the repressor from DNA resulting in transcription.^43^ Thus, TSRs are different from heat shock promoters (HSP) and rely on housekeeping sigma factors such as σA in *B. subtilis*.^44, 45^ TSRs offer a greater dynamic range than HSP and do not necessitate stress conditions for induction.^42, 43^ There is precedent for thermal control of *B. subtilis* with induction of gene expression at low and high temperatures in both native and recombinant systems.^45–50^ Additionally, TSRs have been shown to be controlled previously in Gram-negative organisms with ultrasound to create localized thermal energy for transcriptional control.^27^

The configuration, size and composition of SPIONs have a large effect on MPI performance^53–55^, and magnetothermal heating.^56^ Synomag-D is a commercially available multi-core “nanoflower” particle^57^ and has demonstrated improved MPI performance^58, 59^ as well as high intrinsic power loss under magnetic hyperthermia.^60, 61^ Pairing TSRs with magnetothermal energy using SPIONs for localized heating with AMF can lead to regional transcriptional control as guided by MPI or MRI for new approaches to bacteriotherapy. There are many bacteriotherapy approaches under investigation, and FDA review, for a variety of human cancers.^62–67^ We engineered TSRs^42, 43^ into the model organism, *B subtilis*, towards the development of noninvasive genetic control of a minimally invasive biological therapy (**Figure 1**).

**Figure 1.**
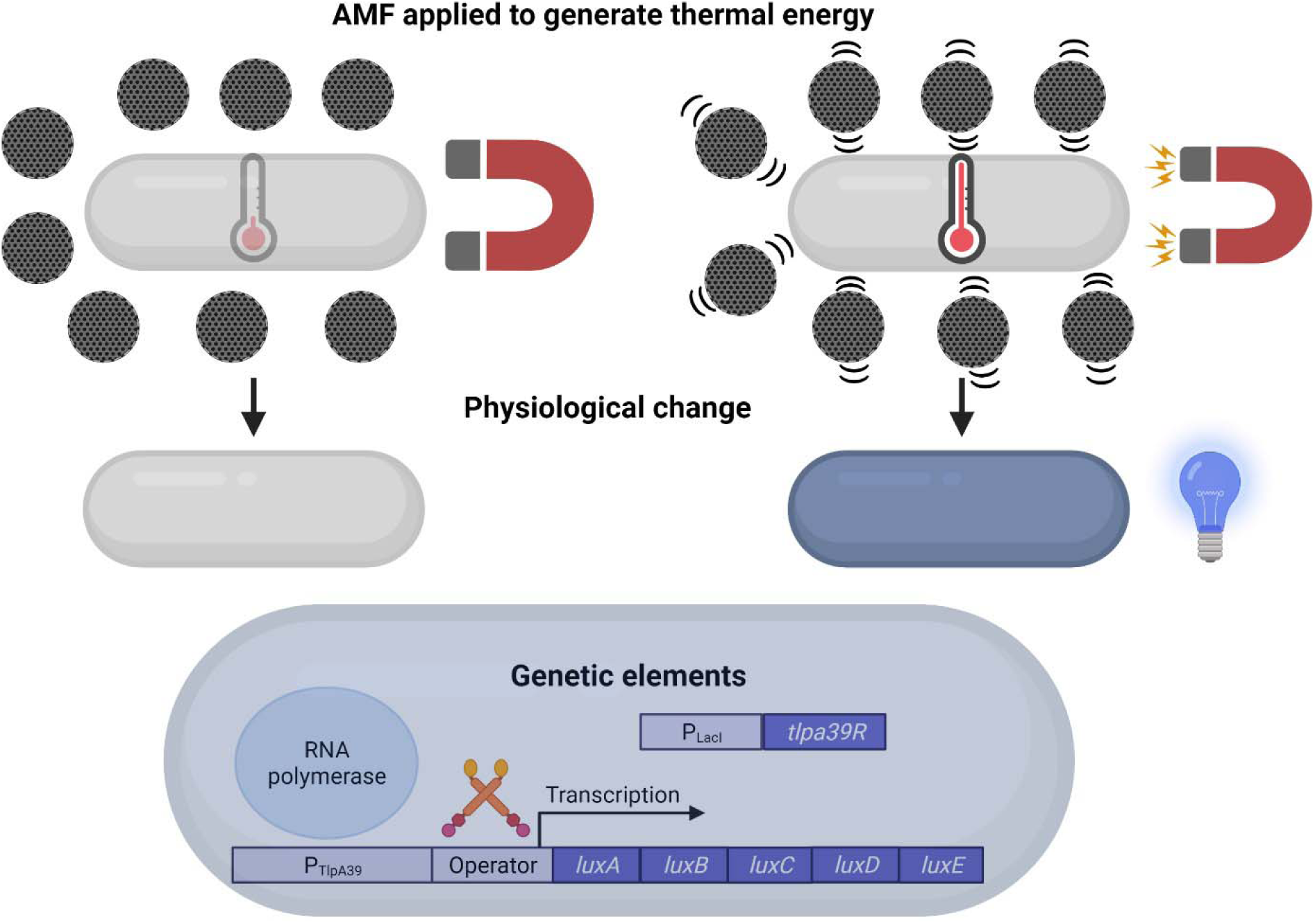
Illustration of *B. subtilis* coated with Synomag-D SPIONs that generate thermal energy upon application of AMF which initiates transcription of *luxA-E* operon from P_TlpA39_.

## Results and Discussion

Magnetic nanoparticles have been used for several biomedical applications and can be further expanded into a measure of control through non-invasive stimuli. Magnetic hyperthermia has been proposed to be used for tumor microenvironment disruption by combining synthetic and biological magnetic nanoparticles with AMF but with limitations.^3, 14, 20, 21, 23^ We investigated the concept of using magnetothermal energy to control a genetic switch in Gram-positive bacteria. This would comprise a modular platform as the basis for developing a variety of potential therapeutics. Directed delivery and targeted activation can improve therapeutic effects and reduce toxicity of bacteriotherapies while imaging can guide development of novel bacteriotherapies by assessing delivery, retention and activation within the target tissue.^68^ Here we use magnetic hyperthermia and imaging to characterize the use of superparamagnetic nanoparticle-coated *B. subtilis* as a new approach for controlling Gram-positive bacterial gene expression with potential use in bacteriotherapies.

A TSR (TlpA39)^42, 43, 69^ was used to control transcription of the the *luxA-E* operon such that luciferase activity (bioluminescence) could be used as a rapid readout for regulation.^70^ This construct demonstrated thermal transcriptional control in *B. subtilis* in response to continuous heating*. B. subtilis* P_TlpA39_ *luxA-E* +*tlpa39R* (+TlpA39R) and *B. subtilis* P_TlpA39_ *luxA-E* -*tlpa39R* (-TlpA39R) were heated continuously in a thermocycler for 12 h at 25°C, 37°C, 39°C or 42°C to test induction of P_TlpA39_. *B. subtilis* +TlpA39R showed a 9.0-fold increase (p<0.0001) in luciferase activity when normalized to OD_600_ from 25°C to 37°C while *B. subtilis* -TlpA39R showed a 0.5-fold increase (p<0.0001; **Figure 2A**). Bioluminescence did not significantly increase when cells were induced at temperatures above 37°C while mRNA levels, as measured by real-time quantitative PCR (RT-qPCR), showed continual increase in P_TlpA39_ activity up to 42°C in *B. subtilis* +TlpA39R (**Figure 2B**). The second gene in the *luxA-E* operon engineered for expression in Gram-positive organisms,^69^ *luxB,* was chosen as the target for RT-qPCR analysis since it encodes for the β subunit of the alkanal monooxygenase enzyme that provides structure for the active conformation of the α subunit of the heterodimeric luciferase.^71^ In the +TlpA39R strain *luxB* levels increased by 1.9-, 3.6-, and 14.6-fold change at 37°C, 39°C and 42°C, respectively. The *tlpa39R* transcript fold change was 1.6, 2.9, and 7.4 at 37°C, 39°C and 42°C, respectively in *B. subtilis* +TlpA39R. *B. subtilis* -TlpA39R showed no significant change in bioluminescence signal and minimal change in P_TlpA39_ activity from mRNA levels as expected from the unregulated promoter (**Figure 2C**). In the -TlpA39R strain *luxB* levels increased by 2.1-, 1.8-, and 1.6-fold change at 37°C, 39°C and 42°C, respectively. The increase in bioluminescence in the -TlpA39R strain from 25°C to 37°C can be attributed to increased activity of the Lux enzymes over those temperatures and more so to a shift in *B. subtilis* metabolism which is consistent throughout the study.^50, 70, 72^ Overall, the results indicate that the P_TlpA39_ promoter and regulator system is functional in *B. subtilis* and able to regulate an operon with a slight temperature shift from what was observed in *Escherichia coli* previously.^42^ This is further demonstrated by the increased levels of *luxB* transcription at increasing temperatures despite the increased levels of *tlpa39R* transcripts indicating more regulator protein available to bind the P_TlpA39_ operator-promoter region as indicated by RT-qPCR. The P_TlpA39_ promoter and regulator system could be further optimized in *B. subtilis* as was done previously in *E. coli*^42^ and *B. subtilis.*^48^ Further optimization by directed mutagenesis^42, 48^ or other measures could improve the P_TlpA39_ genetic switch to have a more stringent on/off state which would be more ideal for *in vivo* studies.

**Figure 2.**
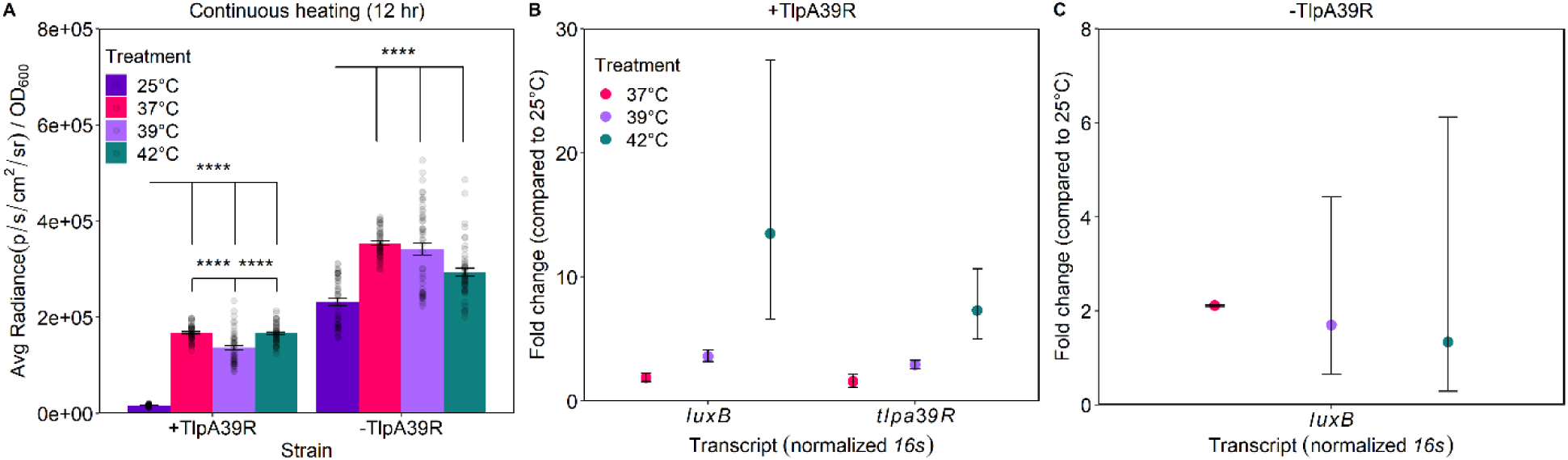
Continuous heating of *B*. *subtilis* +TlpA39R and *B*. *subtilis* -TlpA39R to induce expression of LuxA-E from P_TlpA39_. Error bars are mean ± standard error mean. (A). Transcript levels determined by RT-qPCR for *luxB* and *tlpa39R* from +TlpA39R strain (B) along with *luxB* from -TlpA39R strain (C) at induction temperatures compared to 25°C. RT- qPCR shown as mean with error bars as 95% confidence intervals. Statistics were displayed when comparing to 25°C for both +/- TlpA39R strains and between each increasing temperature in +TlpA39R strain. ****p<0.0001.

To test magnetothermal activation, *B. subtilis* ZB307^73^ (derivative of *B. subtilis* strain 168) was coated SPIONs using plain-dextran, carboxyl or amine-coated Synomag-D^58, 59^; each were assessed for coating efficiency, interactions between SPION and bacteria and magnetothermal heating. Scanning electron microscopy coupled with energy dispersive X-ray spectroscopy (SEM-EDS) was performed and displayed associations with, and retention of, the nanoparticles and *B. subtilis.* All three variations were found surrounding and associating with *B. subtilis* as confirmed by Fe signal from EDS, but with varied consistency of coating observed (**Figure 3**). The plain-dextran and amine-coated evenly covered and associated with *B. subtilis* while the carboxyl-coated appeared to heterogeneously associate with *B. subtilis* in large aggregates (**Figure 3**). Iron signal was absent from a *B. subtilis* sample without SPIONs (**Figure S1)**. SPIONs were not found in the cytoplasm of the *B. subtilis* after coating as shown by transmission electron microscopy (TEM) cross-sections (**Figure S2**). To further investigate the three *B. subtilis* coatings, inductively coupled plasma mass spectrometry (ICP-MS) was performed to measure iron content. There was more iron in the carboxyl-coated samples compared to the plain-dextran and amine-coated, 464.8 and 294.7 times, respectively (**Table S1**). This highlights the disparity between the way the three SPION variations associate with *B. subtilis*.

**Figure 3.**
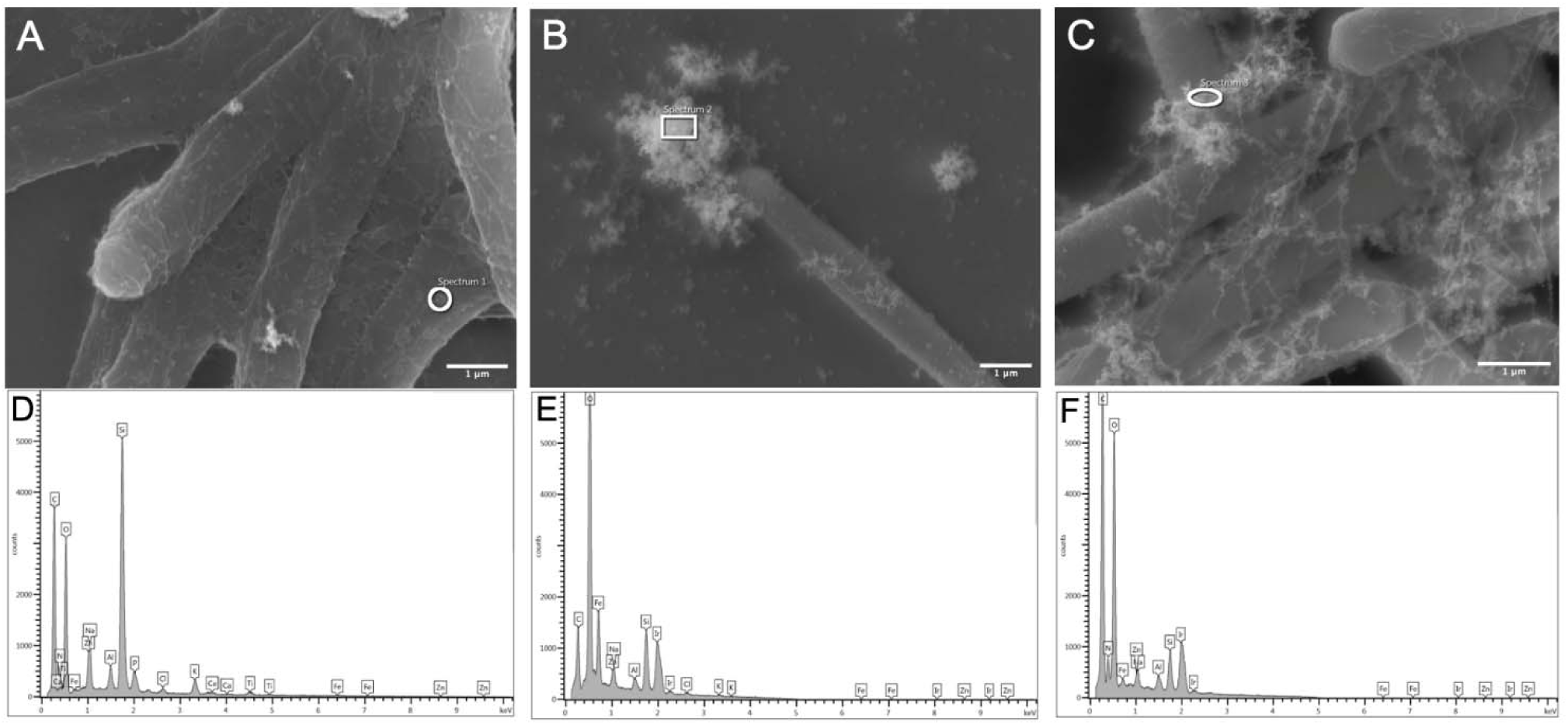
Visualization and elemental analysis of *B. subtilis* and SPION associations. Plain- dextran SPIONs; mag. = 27,000X (A), carboxyl SPIONS; mag. = 23,000X (B) and amine SPIONS; mag. = 33,000X (C) show various associations with *B. subtilis* as observed by scanning electron microscopy. Elemental analysis was performed on each of the samples (D- F) in the regions indicated with white borders to show iron (Fe) signal to identify the SPIONs. Scale bars = 1 µm.

The variations in association and retention of the three types of SPIONs with *B. subtilis* are primarily influenced by electrostatic and dispersive forces between the bacteria and the SPION coatings.^74^ *B. subtilis* has a net negative electrostatic charge and a zeta-potential of – 41 mV when grown at a physiological pH.^75, 76^ Previous studies have demonstrated that with increasing negative zeta-potential, the higher the adhesion potential extends from the bacteria.^76^ Additionally, the DLVO (Derjaguin–Landau–Verwey–Overbeek) theory can be used to explain the potential interaction between a given nanoparticle and bacteria.^74, 76^ The SPIONs used in this study have a net electrostatic charge of negative (plain-dextran),^58, 59^ low negative to neutral charge (carboxyl-coated), or a positive charge (amine-coated) when at physiological pH or pH 6.5 for the amine-coated (MicroMod). Even though the plain-dextran is negatively charged, the difference in zeta potential between *B. subtilis* and the particle was enough to allow for coating similar to previous coatings of *B. subtilis* with gold nanoparticles.^76^ The carboxyl-coated SPION has a high potential for Van der Waals due to its hydroxyl functional groups which contributes to the DLVO theory and increases the aggregation and agglomeration of the nanoparticle in suspension and around *B. subtilis*.^77^ The positive charge of the amine-coated SPION at pH 6.5 promoted association with *B. subtilis* but the pH requirement is a limiting factor for this particle type. Ultimately, the variations in coating between the promising plain-dextran and carboxyl-coated SPIONs at physiological pH were used for downstream applications.

*B. subtilis* viability was assessed by flow cytometry after coating with each of the SPIONs. Two bacterial concentrations, equal to OD_600_ = 1 or 2, were tested while maintaining the same concentration of the Synomag-D variations to determine if a high ratio of iron to *B. subtilis* would cause toxicity. After 2 h of coating, none of the Synomag-D variations at either *B. subtilis* concentrations demonstrated reduction in viability compared to the untreated control and all treatments were significant when compared to the 98°C control for cell death (**Figure 4A, B**). Furthermore, viability was assessed after 12 h of AMF using the plain-dextran particle as it produced the most reproducible heating response from *B. subtilis* +TlpA39R at an OD_600_ = 2 with a 16.0 mT radio frequency (RF) amplitude (data not shown). The *B. subtilis* +TlpA39R strain had slight, but not significant differences in viability compared to the -TlpA39R strain with or without AMF treatment (**Figure 4C**). Overall, the coating and application of AMF did not significantly impact the viability of *B. subtilis*.

**Figure 4.**
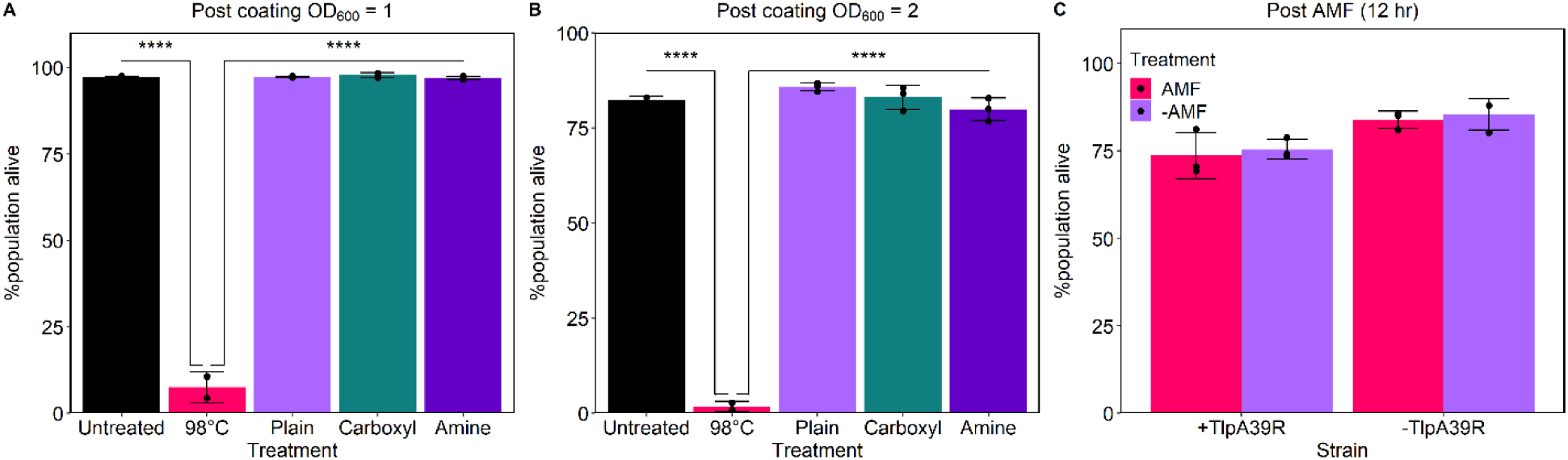
Flow cytometry determining viability of *B. subtilis* after coating with the three- particle variations at two bacterial concentrations (A, B). *B. subtilis* +TlpA39R and *B. subtilis*-TlpA39R were compared in and outside the AMF after 12 h of heating when coated with the plain-dextran particle (C). Error bars are mean ± standard deviation. ****p<0.0001.

To illustrate transcriptional control of potential bacteriotherapies, magnetic hyperthermia using the HYPER system, was applied to *B. subtilis* coated with each of the three variations of the SPION. The growth temperature of the *B. subtilis* for the HYPER experiments was 37°C as opposed to the thermocycler experiments where the growth temperature is 25°C. This was intended to support *in vivo* studies as the core body temperature of mice and humans is approximately 37°C.^78^ Magnetic hyperthermia increased bioluminescent signals in bacteria coated with all particle variations with plain-dextran producing the most reproducible and significant result in the higher bacterial concentration at the max RF amplitude (16.0 mT). The carboxyl-coated SPION caused the highest fold changes in signal compared to the -AMF condition but with the most variability between replicates. The plain-dextran coated *B. subtilis* +TlpA39R showed only a 0.2-fold change (p=0.0456) at the lower concentration when exposed to AMF while at the higher bacterial concentration with a 16.0 mT RF amplitude showed a reproducible 1.4-fold change (p<0.0001; **Figure 5A, D**). Carboxyl-coated *B. subtilis* +TlpA39R showed a 4.6-fold change (p=0.0214) and a 3.4-fold change (p=0.014) in bioluminescence when exposed to AMF at the lower and higher bacterial concentrations respectively but with variability (**Figure 5B, E**). Additionally, the -TlpA39R strain showed a 1.9-fold change (p=0.1689) and a 1.1-fold change (p=0.014) in bioluminescence when exposed to AMF at the lower and higher bacterial concentrations respectively and with high variability. Finally, the amine-coating produced a 0.6-fold increase (p=0.0026) at the lower bacterial concentration in the +TlpA39R strain but showed a small decrease in signal at the higher bacterial concentration when exposed to AMF (p=0.0217; **Figure 5C, F**). Due to the plain-dextran coating producing the most significant and reproducible result at the higher bacterial concentration, this condition was chosen for transcript measurements. There was a 1.2-fold increase in *luxB* levels even with increasing *tlpa39R* levels (1.7-fold change) in the +TlpA39R strain (**Figure 5G**) and 1.4-fold increase in *luxB* in the -TlpA39R strain after AMF exposure (**Figure 5H**).

**Figure 5.**
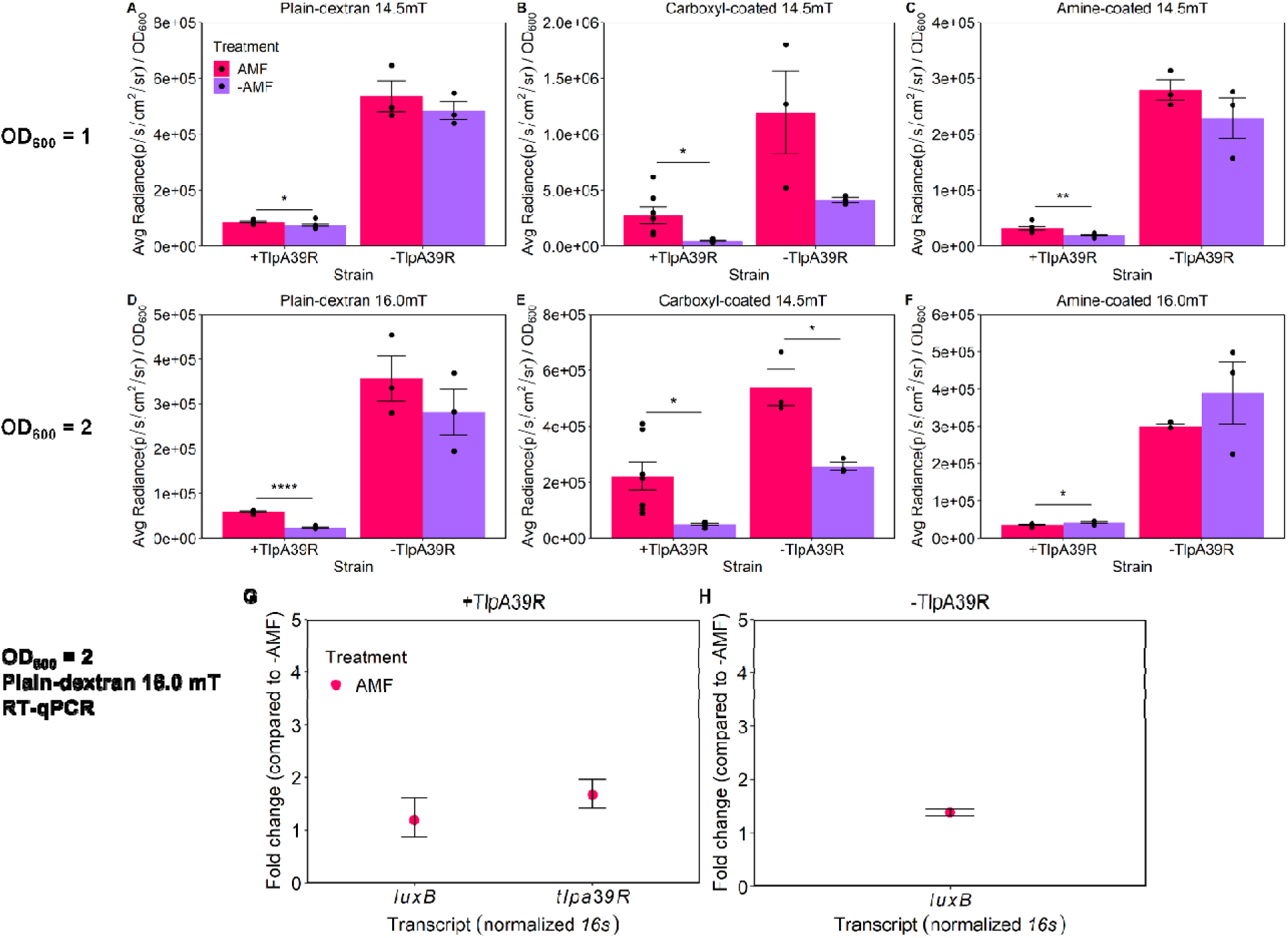
Magnetic hyperthermia increasing bioluminescent signal (Avg. Radiance) using the HYPER Theranostic Hyperthermia System. *B. subtilis* +TlpA39R and *B. subtilis* -TlpA39R were compared in and outside the AMF with the three Synomag-D coating variations at OD_600_ = 1 (A-C) or 2 (D-F). Error bars are mean ± standard error mean. RT-qPCR was used to determine transcript levels of the two strains and compare AMF to -AMF(G-H). RT-qPCR shown as mean with error bars as 95% confidence intervals. *p<0.05, **p<0.01, ****p<0.0001.

The SEM-EDS and TEM (**Figure 3, S2**) provide some explanation for the results seen following magnetic hyperthermia. The plain-dextran and amine-coated SPIONs evenly coated *B. subtilis* while the carboxyl-coated SPIONs formed large aggregates that indicated potentially more iron around *B. subtilis* but with differences between bacteria in the sample. Therefore, AMF could result in greater thermal energy being delivered to *B. subtilis* through the carboxyl-coated SPION than with the plain-dextran or amine-coated SPION, but with higher variability due to less reproducible associations with the bacteria. The HYPER parameters were chosen based on several preliminary experiments that optimized RF amplitude for each particle at each bacterial concentration then the best conditions for each particle were performed with maximum biological replicates that could be placed inside the HYPER system (**Figure 5**). Thermal probes measuring the temperature of the culture medium showed that the carboxyl-coated SPION was the only particle that increased culture medium temperature (+3°C) when exposed to AMF (**Figure S3**). This was supported by the electron microscopy indicating more free iron throughout the media in addition to the aggregates associated with the bacteria **(Figure 3B, S2E).** The plain-dextran and amine-coated SPIONs did not increase temperature but still induced P_TlpA39_ indicating potential direct thermal energy transfer to *B. subtilis*. Classical heat transfer theory based on Fourier’s law of thermal conduction could explain this phenomenon at the micrometer scale taken together with coating observed under SEM-EDS but thermal confinement to *B. subtilis* is unlikely.^79^ Explaining the observed thermal energy transfer phenomenon by Fourier’s law is also supported by the observed differences in heating between the three particle variations. The carboxyl-coating caused the largest fold change, though variable, and also increased the culture medium temperature which would be consistent with the law of thermal conduction.^79, 80^ Accordingly, the other two particle variations were diffusing thermal energy that did not cause a detectable culture medium temperature change but could have still caused the biological response from *B. subtilis* especially when comparing the +/- TlpA39R strains. We chose the plain-dextran SPION for 1 h thermal induction and *in vivo* studies because of the reproducibility of heating, even coating of *B. subtilis* to maximize retention and less thermal energy transfer throughout the culture medium which could translate to less damage to surrounding tissue *in vivo.* In future studies, the SPION of choice should be determined based on desired effects as the varied particle characteristics could have different advantages in other scenarios.

For small animal *in vivo* applications, anesthesia for times >1 h can cause negative impacts on animal health.^81, 82^ Therefore, reducing AMF application time to around 1 h was necessary for demonstration of translatability of this approach. Increases in bioluminescent signals and *luxB* levels were seen after 1 h of continuous heating in a thermocycler (**Figure 6A, B**). *B. subtilis* +TlpA39R had a 11.1-fold increase (p<0.0001) in luciferase activity when normalized to OD_600_ from 25°C to 37°C while *B. subtilis* -TlpA39R showed a 1.3-fold increase (p<0.0001). When increasing the temperature from 37°C to 39°C and from 39°C to 42°C in the regulated strain (+TlpA39R) there was a 0.5-fold (p<0.0001) and 0.1-fold change (p<0.0001) between each step up, respectively, whereas the –TlpA39R strain had a negative fold change when comparing bioluminescent signal between 37°C to 39°C (-0.02-fold;p=0.9061) and 39°C to 42° (-0.3- fold;p<0.0001). In the +TlpA39R strain *luxB* transcript levels increased by 0.9-, 0.9-, and 1.8- fold change at 37°C, 39°C and 42°C, respectively, indicating induction between 39°C and 42°C. The *tlpa39R* transcript fold change was 0.8, 0.9, and 0.9 at 37°C, 39°C and 42°C, respectively in *B. subtilis* +TlpA39R showing consistent levels as temperature increased. The *B. subtilis* - TlpA39R showed similar changes to +TlpA39R in *luxB* levels at 37°C and 39°C (0.9- and 0.9- fold, respectively), but showed a lesser fold change of 0.4 at 42°C compared to the regulated strain (**Figure 6C**). This indicates that there is some temperature dependent induction of *luxB* in the +TlpA39R strain after 1 h of continuous heating. However, AMF application for 1 h only increased bioluminescent signal by 0.02-fold in the +TlpA39R strain and a minimal 1.0-fold increase (doubling) in *luxB* transcripts which was similar to the –TlpA39R *luxB* mean increase of 1.5 (**Figure 6D-F**). As was shown in the comparison of 12 h of continuous heating to AMF (**Figure 2, 5**), magnetic hyperthermia does not induce the TSRs to the same degree as continuous heating. Accordingly, results obtained after 1 h of heating indicate that continuous heating over this limited time can significantly increase the output of the reporter, which demonstrates the potential for *in vivo* use. However, it is likely that the pulse sequence of magnetic hyperthermia used here would need to be improved to maximize potential for *in vivo* applications. An immediate change to the current process, that could enhance the magnetic hyperthermia, is increasing the RF amplitude beyond the limitations of the HYPER system (>16.0 mT). However, as an increase in RF amplitude will result in an increase in specific absorption rate (SAR)^83^, this would have to be further studied to prevent any biological effects. The plain-dextran particle used here to coat *B. subtilis* is promising and shows potential for enhanced thermal energy transfer from a stronger AMF. Alternatively, other SPIONs could be investigated to further enhance magnetic hyperthermia response in *B. subtilis.* Various SPIONs have been modified to improve magnetic hyperthermia properties^84–87^ and these variations should be investigated for efficient coating of *B. subtilis* and improved magnetic hyperthermia after exposure to AMF.

**Figure 6.**
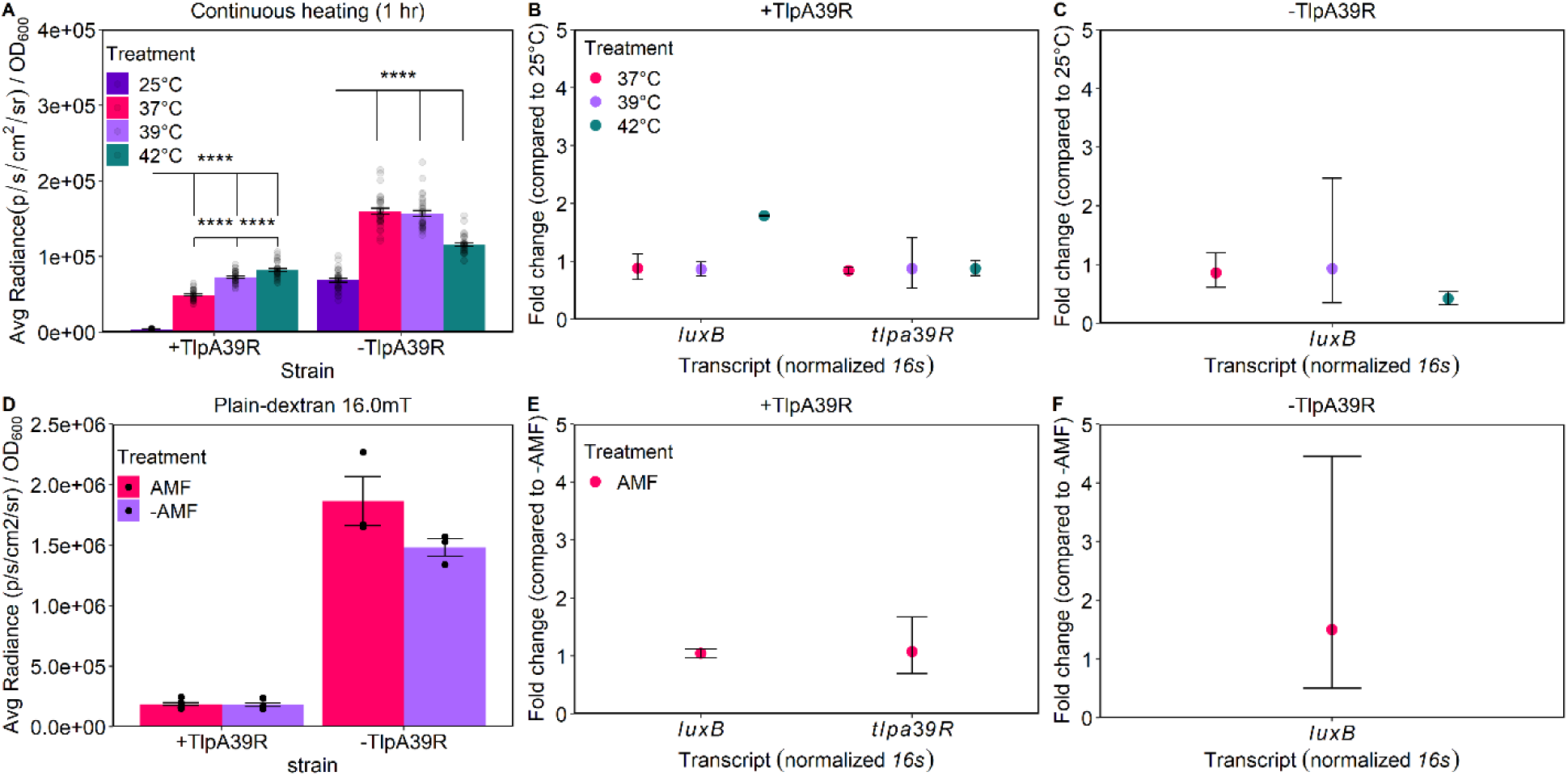
One-hour thermal inductions using continuous heating and magnetic hyperthermia with reporter gene activity (Avg. Radiance) and transcript levels measured (A-C). *B. subtilis* +TlpA39R and *B. subtilis* -TlpA39R were compared in and outside the AMF (D-F). Error bars are mean ± standard error mean. RT-qPCR shown as mean with 95% confidence intervals. Statistics were displayed when comparing to 25°C for both +/- TlpA39R strains and between each increasing temperature in +TlpA39R strain for continuous heating.

MPI was performed to quantify iron content in each sample, and these values were compared to those identified by ICP-MS (**Table S1**). Samples containing 1x10^8^ *B. subtilis* coated with the three Synomag-D coatings were resuspended in a volume relevant to intramuscular (IM) injections (25 µL). Only the carboxyl-coated *B. subtilis* could be detected in these conditions (**Figure 7B**), with iron concentration at 384.8 ppm. The plain-dextran could not be detected when the 25 µL samples were imaged using MPI neither *in vitro* (**Figure 7A**) nor *in vivo* (**Figure S4**). When the plain-dextran samples were pooled to a total volume of 100 µL, MPI signals were detected (**Figure 7C** **inset)** and iron quantified was 0.5 ppm, or 13.6 ng per 25 µL sample injected *in vivo* (**Figure 7D**). The amine-coated sample was not detectable in a 25 µL sample volume (**Figure 7C**) and was not pursued further due to the poor AMF response observed previously (**Figure 5C,F**). MPI quantification showed that the carboxyl-coated SPION sample was 707.3 times that of the plain-dextran SPION (**Figure 7D**).

**Figure 7.**
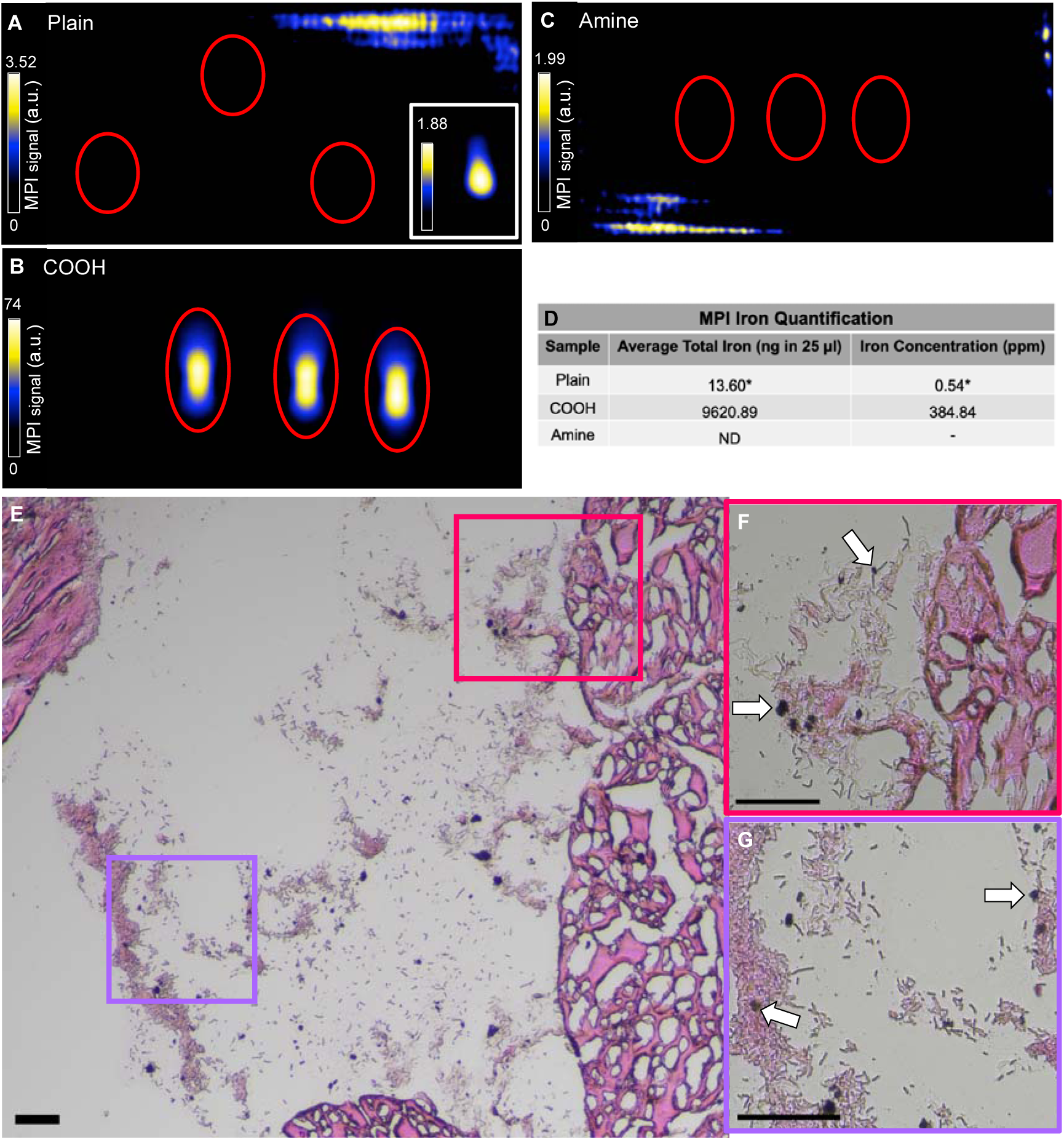
MPI and histological analysis. *B. subtilis* +TlpA39R coated with the three SPION variations were analyzed via MPI in triplicate. MPI signals could not be detected in the plain- dextran sample in a 25 µL volume (A). Inset shows a pooled volume of 100 µL plain-dextran sample, adjusted to visualize MPI signals. Carboxyl-coated samples showed signal (B) while the amine-coated samples (C) were not detected in 25 µL volumes. MPI scale bars are individual for each condition and represent the full dynamic range of the image. Iron content was quantified using MPI data (D). *Quantification of the plain-dextran sample was performed on an 100 µL pellet and then calculated for a 25 µL volume. Sectioned tissue stained with a modified Gram stain followed Perls’ Prussian Blue (PPB); magnification = 100X (E) and zoomed in regions indicated by magenta (F) and purple (G) boxes with magnification = 400X. White arrows indicate PPB-stained iron. a.u. = arbitrary units; ND = not detected. Scale bars = 50 µm.

A murine model of IM thigh injections was paired with the HYPER system and histology to assess iron association with the bacteria and potential changes in bioluminescence. Bacteria coated with SPIONs were prepared and imaged for bioluminescence quantification pre-injection, immediately post-injection, and 1 h post-treatment (+/-AMF). The bioluminescence levels decreased 4.1-fold and 8.0-fold from pre-injection to post-injection in +AMF and -AMF treatments, respectively, but the variance was high between replicates so there was no significance (p=0.4841; p=0.3446; **Figure S5**). The change in bioluminescence before and after treatment was negligible with a 0.43-fold decrease and a 0.56-fold increase +AMF and -AMF, respectively (p=0.4813; p=0.4760; **Figure S5**). Histology using a modified Gram stain confirmed presence of *B. subtilis* within sectioned intramuscular tissue after 1 h treatments. Further, consecutive staining with a Perls’ Prussian Blue protocol revealed *B. subtilis* and iron staining in the same location within the tissue (**Figure 7E-G**). The white arrows (**Figure 7F-G**) indicate the presence of iron due to the insoluble Prussian blue pigment which is formed after the potassium ferrocyanide reagent reacts with ferric iron in the sample.^88^ These images suggest there is association and retention of the SPION with the bacteria after injection *in vivo*, but could not be confirmed with the optical microscopy technique utilized. The use of the consecutive staining scheme which included multiple counterstains and decolorizing steps led to an atypical Gram stain result for *B. subtilis*. A sequential tissue section was stained using only the modified Gram stain and the standard purple rods of *B. subtilis* were observed adjacent to the muscle tissue stained yellow from the alcoholic saffron (**Figure S6A-C**). The modified Gram stain further supported the finding of the association of the *B. subtilis* and plain-dextran SPION *in vivo* by showing a typical Gram stain result for *B. subtilis* in comparison to the consecutive staining (**Figure 7** **E-G**). Hematoxylin and eosin staining performed on adjacent tissue sections showed eosin-stained (pink), longitudinal quadriceps muscle fibers (**Figure S6D**) confirming samples were injected intramuscularly.

Perls’ Prussian Blue staining^89^ and modified Gram stain^90^ demonstrated the presence of iron and *B. subtilis* at the same location, which provides the opportunity to utilize magnetic hyperthermia to control *B. subtilis* transcription *in vivo.* Further tuning of the genetic elements to the *B. subtilis* and characterizing the interaction of improved particles for magnetic hyperthermia with *B. subtilis* would enhance further *in vivo* studies. SPIONs can be coated with polymers, small molecules, lipids and composites to increase stability, water solubility and biocompatibility.^85^ For example, Fe_3_O_4_-oleic acid-Na-oleate nanoparticles^87^ increased stability in a transplanted carcinoma model and polycaprolactone-coated superparamagnetic iron oxide nanoparticles synthesized with a micellular conformation to increase cytocompatibility and thermosensitivity as a cancer therapy.^85^ Additionally, increasing RF amplitude and amount of iron associating with the bacteria could improve heating along with imaging properties *in vivo*.

Yet, increases in bioluminescence were observed after AMF treatment with only ∼1 ppm of Fe in the plain-dextran coated condition *in vitro*. This reduced the amount of Fe that is delivered for other magnetic hyperthermia applications such as for tumor ablation from 1 mg/cm^3^ to 13.6 ng/cm^3^ that was used in our study.^91^ Accordingly, the bacteria can be used as a carrying mechanism for and a responsive mechanism to SPIONs where minimal SPIONs are needed to produce a desired therapeutic outcome through controlling bacteriotherapies. Alternatively, manganese-doped magnetic nanoclusters have been studied for glioblastoma therapy as a nanoparticle that has complementary functionalities and can utilize photothermal and magnetic hyperthermia treatments.^86^ Additionally, other heating mechanisms could be used for magnetic hyperthermia such as ultrasound which was shown previously.^42, 52, 92^

## Conclusions

The introduction of the TlpA39 regulatory system into *B. subtilis* demonstrated that a temperature-sensitive repressor optimized in a Gram-negative organism can be utilized in a Gram-positive organism to drive controlled transcription of the *luxA-E* operon by continuous or magnetothermal heating. After coating the bacteria with three SPION variations (plain-dextran, carboxyl, amine), SEM-EDS confirmed that the plain-dextran and amine SPIONs coated the *B. subtilis* in an even, thin coating compared to the carboxyl-coated particle that formed large aggregates that heterogeneously associated with *B. subtilis* without impacting viability. Both thermocycler heating and magnetic hyperthermia by the HYPER system created substantial increase in bioluminescent output over 12 h but only continuous heating showed a significant increase in 1 h. This indicated that thermal induction is possible to measure in timeframes relevant to *in vivo* studies with small animals, but the genetic elements need to be tuned for *B. subtilis* to enhance the switch between off/on states after 1 h of stimulus and other nanoparticles should be tested to improve magnetic hyperthermia. Yet, the SPIONs do appear to maintain association with *B. subtilis* after injection *in vivo* indicating that magnetic hyperthermia can be used to control bacterial transcription if longer heating timeframes can be used (*e.g*., larger animal studies) or enhancements mentioned above are performed.

## Materials and Methods

### Data and code availability

All raw data, *Bacillus subtilis* constructs and R scripts will be made available upon request by the corresponding author. Plasmids used to produce *B. subtilis* constructs will be submitted to Addgene after manuscript publication. All R scripts were written with established packages.

### Bacterial growth conditions

*B. subtilis* constructs were grown in Luria-Bertani Miller broth (LB) with spectinomycin (100 µg/mL). The overnight cultures were grown for 16 h at 37°C and 250 RPM unless otherwise specified.

### B. subtilis constructs

The thermal response elements originated from pTlpA39-Wasabi (Addgene plasmid # 86116; http://n2t.net/addgene:86116; RRID:Addgene_86116).^42^ The TlpA39 promoter and regulator (driven by the LacI promoter) were cloned into the pDR111 plasmid to replace the Phyper-spank promoter and LacI regulator using Gibson assembly.^93^ Accordingly, the *luxA-E* operon was inserted into in the NheI restriction site of the pDR111 backbone by the seamless ligation cloning extract (SLiCE) method^94^ to create the new pDR111 P_LacI_ *tlpa39R* P_TlpA39_ *luxA-E. coli* construct. Three strains were created: empty vector (pDR111 backbone only), experimental P_TlpA39_ repressed strain (pDR111 P_LacI_ *tlpa39R* P_TlpA39_ *luxA-E*), and P_TlpA_ constitutive strain without the repressor (pDR111 P_TlpA39_ *luxA-E*). Constructs were inserted into the genome of *B. subtilis* at the *amyE* locus using a homologous recombination plasmid (pDR111^95^, a gift from Dr. Lee Kroos). The pDR111 plasmid was transformed into *B. subtilis* using a natural competence protocol and constructs were selected for by spectinomycin then confirmed by PCR amplification out of the genome.^96^ Three *B. subtilis* strains were created: containing the empty vector, the vector with the experimental P_TlpA39_ repressed strain (P_TlpA39_ *luxA-E* + *tlpa39R*), and the P_TlpA_ constitutive strain without the repressor (P_TlpA39_ *luxA-E* - *tlpa39R*). All constructs were confirmed by PCR, restriction enzyme digestion, functional assays (when applicable), and Sanger sequencing (Azenta Life Sciences). A list of all oligos used is available in supplementary information (**Table S2**).

### Iron coating of *B. subtilis*

Synomag-D particles possess a maghemite (γ-Fe_2_O_3_) core of nanoflower-shaped nanocrystallites with a dextran shell and a hydrodynamic particle diameter of 50 nm.^59^ We utilized the plain dextran shell nanoparticle, a variation coated with carboxyl groups (carboxyl-coated) and a variation coated with amine groups (amine-coated) (MicroMod; #104-00-501, #103-02-501, #104-01-501). *B. subtilis* was incubated with plain-dextran or carboxyl-coated Synomag-D (200 µg/mL) in LB broth (pH = 7) or in LB broth (pH = 6.5) for the amine-coated Synomag-D (200 µg/mL) for 2 h at 37°C and 250 RPM after being normalized to OD_600_ = 1 or 2 in 1 mL. Coated *B. subtilis* was spun down at 10,000 x g for 2 min and washed with PBS (pH = 7.4) for plain- dextran and carboxyl-coated Synomag-D or with PBS (pH = 6.5) for amine-coated Synomag-D. The cultures were then resuspended in 100 µL of LB broth with appropriate pH mentioned above for use in HYPER or 250µL of PBS (pH appropriate) for MPI and *in vivo* experiments.

### Scanning electron microscopy and elemental analysis

Five hundred microliters of coated *B. subtilis* suspended in growth media was mixed with an equal volume of 4% glutaraldehyde in 0.1M sodium phosphate buffer, pH 7.4. Fixation was allowed to proceed for 30 min at room temperature. Twelve-millimeter round glass coverslips were floated on one drop of 1% poly-L-lysine (Sigma Aldrich P1399) each and allowed to stand for 10 min. The coverslips were removed and gently washed with HPLC-grade water. One drop of fixed sample was placed on the now coated side of the coverslip and allowed to settle for 10 m. After sample addition the coverslip was gently washed with HPLC-grade water and placed in a graded ethanol series (25%, 50%, 75%, 95%) for 10m each with three 10m changes in 100% ethanol.^97^

Coverslips with samples were then critical point dried in a Leica Microsystems model EM CPD300 critical point dryer (Leica Microsystems, Vienna, Austria) using carbon dioxide as the transitional fluid. Coverslips were then mounted on aluminum stubs using System Three Quick Cure 5 epoxy glue (System Three Resins, Inc., Aubur, WA) and carbon conductive paint (Structure Probe, Inc. 05006-AB) was added in a thin line for grounding. Samples were coated with iridium (2.7 - 5.5 nm thickness) in a Quorum Technologies/Electron Microscopy Sciences Q150T turbo pumped sputter coater (Quorum Technologies, Laughton, East Sussex, England BN8 6BN) purged with argon gas.

Samples were examined in a JEOL 7500F (field emission emitter) scanning electron microscope (JEOL Ltd., Tokyo, Japan) and energy dispersive X-ray spectroscopy (elemental analysis) was performed using an Oxford Instruments AZtec system (Oxford Instruments, High Wycomb, Bucks, England), software version 3.1 using a 150mm^2^ Silicon Drift Detector (JEOL 7500F SEM) and an ultra-thin window. Images were analyzed using Fiji (ImageJ, version 2.0.0- rc-69/1.52i).

### Transmission electron microscopy

Transmission Electron Microscopy (TEM; JEM-1400Flash, JEOL, MA USA) was used to confirm external associations of SPIONs with *B. subtilis*. Pelleted samples were fixed in 2.5% EM-grade glutaraldehyde for 5 min, washed with 0.1M phosphate buffer, and post-fixed with 1% osmium tetroxide in 0.1M phosphate buffer. After fixation, samples were dehydrated in a gradient series of acetone and infiltrated and embedded in Spurr’s resin. Seventy nanometer thin sections were obtained with a Power Tome Ultramicrotome (RMC, Boeckeler Instruments. Tucson, AZ), floated onto 200-mesh, carbon-coated formvar copper grids. Images were taken with JEOL 1400-Flash Transmission Electron Microscope (Japan Electron Optics Laboratory, Japan). Images were analyzed using Fiji (ImageJ, version 2.0.0-rc-69/1.52i).

### *In vitro* imaging

Plain, carboxyl or amine Synomag-D coated *B. subtilis* were imaged in triplicates (1x10^8^ cells per sample in 25 µL PBS) using the Momentum MPI scanner (Magnetic Insight Inc, CA, USA). Plain Synomag-D coated *B. subtilis* were combined to a total of 4x10^8^ cells in 100 µL PBS for detection. Images were acquired using a 2D projection scan with default (5.7 T/m gradient) or high sensitivity (3 T/m gradient) settings, rf amplitude (16.5 mT x-channel, 17 mT z-channel) and 45 kHz excitation with a field of view (FOV) = 12 x 6 cm, 1 average and acquisition time of ∼1 minute.

Bioluminescence was measured on the *in vivo* imaging system (IVIS, PerkinElmer) with auto-exposure settings (time = 2-40 sec, binning = medium, f/stop = 1, emission filter = open). Average radiance (p/sec/cm2/sr) was normalized to bacterial growth using optical density measured as absorbance at 600 nm (OD_600_) on a plate reader (Spectra Max 3, Molecular Devices, San Jose, CA, USA). Bioluminescent signals were quantified using the 8x12 grid ROI for all wells (*in vitro* thermocycler induction) and ellipse ROIs with standardized area for all tubes (*in vitro*) to calculate average radiance (p/sec/cm2/sr) using Living Image software (PerkinElmer, Version 4.5.2).

### Flow cytometry determination of *B. subtilis* viability

The effects of coating and heating on *B. subtilis* viability was assessed using flow cytometry. *B. subtilis* was coated as described above with all three nanoparticle variations (+/- AMF). Following treatment cells were resuspended in 100 µL of 150 mM NaCl and stained using a viability/cytotoxicity assay kit for live and dead bacteria (Biotium, #30027) according to the manufacturers protocol. Following staining cells were collected by centrifugation and washed once with flow buffer (1X PBS, 0.5% bovine serum albumin) followed by fixation with 4% paraformaldehyde for 10 minutes. Cells were then resuspended in 100 µL flow buffer for analysis using the Cytek Aurora flow cytometer. Unstained dead (heat treated; 98°C), live (uncoated; untreated) and live (coated; plain, carboxyl, amine) plus single stained DMAO (live/dead, FITC) and Ethidium Homodimer III (EthD-III; dead, Cy3) were used as controls. EthD-III dead cells were gated on the DMAO+ cell population. Data were analyzed using FCS express software (De Novo Software, CA, USA; version 7.12.0005). A one-way ANOVA was used to determine any significance between treatments’ potential impact on viability. The data presented herein were obtained using instrumentation in the MSU Flow Cytometry Core Facility. The facility is funded in part through the financial support of Michigan State University’s Office of Research & Innovation, College of Osteopathic Medicine, and College of Human Medicine.

### RNA extraction and RT-qPCR

Technical replicates from thermal inductions were pooled for RNA extractions. *B. subtilis* was lysed using LETS buffer (100mM LiCl, 10mM EDTA, 10mM Tris pH 7.8, 1% SDS) and bead beating (0.1mm zirconium beads, 3 cycles of 60 sec at max speed). Total RNA was extracted using RNeasy miniprep kit (QIAGEN). Samples were cleaned and made into cDNA with QuantiTect Reverse Transcription kit (QIAGEN). The resulting cDNA was diluted 1:20 in RNAse free water for qPCR. QuantiTect SYBR Green PCR kit (QIAGEN) was used to prepare 20 µL reactions according to instructions. Primers for *luxB*, *16s*, and *tlpa39R* were created using NCBI Primer BLAST and used for all samples (**Table S2**). No-template controls of RNase-free water were run in triplicate for each primer set. Reactions were run in triplicate for each sample. Data was screened for validity using melting curves and then analyzed for relative quantification using the 2^-ΔΔCt^ method.^98^ The expression levels for *luxB* and *tlpa39R* was calculated relative to the *16s* rRNA housekeeping gene and the experimental groups (37, 39, 42°C). Confidence intervals of 95% were calculated using the mean and standard deviation for all cycle threshold values of a given sample (n=6) and then converting to fold change using the above 2^-ΔΔCt^ method.

### *In vitro* thermocycler induction

Cultures of *B. subtilis* were grown in LB for 36 h at 25°C and 250 RPM under spectinomycin (100 µg/mL) selection. These cultures were then diluted to optical density of 0.1 at 600 nm (OD_600_) in LB with appropriate antibiotic and grown until they reached OD_600_ of 0.25. Twenty-five microliters of the samples were aliquoted into 96-well PCR plates and sealed. Thermal inductions were carried out in Biorad C-100 thermocyclers at 25, 37, 39, and 42°C for either 12 h or 1 h. After thermal induction, the samples were diluted 1:4 in LB and 90 µL was transferred to a 96-well, black Costar plate. The OD_600_ and bioluminescence output was measured. Controls for the measurements were: growth media only, empty vector strain, P_TlpA39_ constitutive strain without the repressor, and experimental P_TlpA39_ repressed strain. After bioluminescent imaging and optical density measurements (method above), a one-way ANOVA with Tukey’s post-hoc was used to determine any significance between temperatures within a strain.

### HYPER Theranostic Hyperthermia System

Magnetic hyperthermia was performed using the HYPER Theranostic Hyperthermia System (Magnetic Insight). Magnetothermal heating is localized using a Field Free Point (FFP) to direct radiofrequency (RF) energy. HYPER was programmed to apply AMF to the coated *B. subtilis* strains by using a 0.66 T/m magnetic field gradient strength, a RF amplitude of 14.5 or 16.0 mT, 350 kHz excitation and a RF amplitude application time of 60 seconds with a 1 second cool down time. This programmed AMF cycle would be repeated such that the AMF was applied for the desired total run time of either 1 or 12 h. Optimal parameters for each particle were determined when bacteria were normalized to OD_600_ = 1 or 2. For each run the following strains and replicates were included: *B. subtilis* P_TlpA39_ *luxA-E* +TlpA39R coated with one of the three variations of Synomag-D were divided into PCR tubes in two 50 µL aliquots where one aliquot would be placed in the AMF (+AMF) and outside the AMF (-AMF) in biological replicates (n=7). The same process was repeated for the *B. subtilis* P_TlpA39_ *luxA-E* -TlpA39R strain (n=3) to be run in same conditions with +TlpA39R strain. Optimal HYPER parameters for each variation of particle mentioned above with bacteria normalized to OD_600_ = 2 were utilized to determine temperature increase in LB during the heating of *B. subtilis* with the three variations of coating. Fiber-optic temperature probes (Weidmann-Optocon, standard TS2 probes) were placed into the LB throughout the 12 h of heating to track temperature through the HYPER software. Temperature readings were recorded every AMF application cycle (60 readings per cycle were treated as technical replicates) with reads at each 30 min time point plotted for visualization. Unpaired Student or Welch’s t-test was used to determine statistical significance between samples with and without AMF treatment.

### *In vivo* magnetothermal heating and imaging

Female BALB/c mice (6-8 weeks; Jackson Laboratories USA) were obtained and cared for in accordance with the standards of Michigan State University Institutional Animal Care and Use Committee. *B. subtilis* were coated with plain Synomag-D as described above. Mice (n=6) were anesthetized with isoflurane administered at 2% in oxygen followed by hair removal on each thigh using a depilatory. An intramuscular (IM) injection of 1x10^8^ iron-coated bacteria in 25 µL PBS was performed into the left thigh followed by BLI (IVIS Spectrum; post-injection timepoint) using auto-exposure settings (time = 30-120 sec, binning = medium, f/stop = 1, emission filter = open). One mouse was imaged using MPI using the default setting, as described above. No signal was detected and no further mice were imaged by MPI. Following imaging, mice were either placed into the HYPER system for magnetothermal heating (+AMF; 16 mT for 1 hour, n = 3) or maintained at room temperature in cage (-AMF, n = 3). BLI was performed as above, after AMF application, or 1 h for mice which were not subjected to AMF (post-treatment timepoint). After the final imaging time point mice were sacrificed and thigh muscle from the IM injected side and the contralateral not injected side were excised followed by sectioning for histological staining and microscopy (see below for detailed methods). Two-way repeated measures ANOVA and Tukey’s post-hoc was used to determine statistical significance between AMF treatments and timepoints for bioluminescence.

### Histological Analysis

Thigh muscle samples were fixed in 4% paraformaldehyde for 24 h followed by cryopreservation through serial submersion in graded sucrose solutions (10%, 20% and 30%). Samples were then frozen in optimal cutting temperature compound (Fisher HealthCare, USA). Tissues were sectioned using a cryostat (6 µm sections). Sections were stained with a modified Gram stain as described by Becerra *et al*., 2016,^90^ followed by Perls’ Prussian Blue (PPB)^89^ on the same sections for detection of bacteria and detection of ferric iron. CitriSolv (Decon Labs, Inc., King of Prussia, PA, USA; Cat. #1601) was used as a safe alternative for xylene in the final step of the modified Gram stain. Eosin was used as a counterstain in the Perls’ Prussian Blue protocol. Sequential sections of the tissue were stained with the modified Gram stain only and hematoxylin and eosin staining only to verify Gram status of the *B. subtilis* and confirm intramuscular injections, respectively. Microscopy was performed on the sections using a Nikon Eclipse Ci microscope equipped with a Nikon DS-Fi3 camera (Nikon, Tokyo, Japan)) for color image acquisition and NIS elements BR 5.21.02 software (Nikon). Microscopy images were prepared using the auto-white feature on NIS elements and Fiji (ImageJ, version 2.0.0-rc- 69/1.52i).

### Image analysis

Living Image software (PerkinElmer, Version 4.5.2) was used to quantify bioluminescent signals. An 8x12 grid region of interest (ROI) was used for 96-well plates (*in vitro* thermocycler induction) or ellipse ROIs with standardized area for all tubes (*in vitro*) or on the injection site of the mouse thigh (*in vivo*) to calculate average radiance (p/sec/cm2/sr).

MPI data sets were visualized and analyzed utilizing Horos imaging software (Horos is a free and open-source code software program that is distributed free of charge under the LGPL license at Horosproject.org and sponsored by Nimble Co LLC d/b/a Purview in Annapolis, MD USA).

Fixed ROIs were used to identify all samples and total MPI signal was determined (area x mean signal). Calibration standard curves were created by imaging different amounts of iron and plotting signal (y) versus iron content (x) with the y-intercept (b) set to zero. The slope (m) of the data was found using a simple linear regression and quantification of iron content was calculated using the trendline equation (y=mx+b). Standard curves were created using matched imaging parameters (default or high sensitivity) dependent on the data set being analyzed.

### Inductively Coupled Plasma-Mass Spectrometry (ICP–MS)

After *B. subtilis +* TlpA39R was coated with the three SPION variations at OD_600_ = 2 then the cells were pelleted (centrifugation at 10,000xg) and resuspended in phosphate buffered saline (PBS; pH 7.4). Three technical replicates of the coating procedure were pooled in a final volume of 750µL PBS. One sample of untreated *B. subtilis +* TlpA39R (OD_600_ = 2) from the coating process (no SPION control) was also prepared and resuspended in 250µL PBS. The cells were digested in concentrated nitric acid (J.T. Baker, USA; 69-70%) overnight, and diluted 25-fold with a solution containing 0.5% EDTA and Triton X-100, 1% ammonium hydroxide, 2% butanol, 5 ppb of scandium, and 7.5 ppb of rhodium, indium, and bismuth as internal standards (Inorganic Ventures, VA, USA). The samples were analyzed on an Agilent 7900 ICP mass spectrometer (Agilent, CA, USA). Elemental concentrations were calibrated using a 5-point linear curve of the analyte-internal standard response ratio. Bovine liver (National Institute of Standards and Technology, MD, USA) was used as a control.

### Statistical analysis and visualization

Statistical analyses were performed using Prism software (9.2.0, GraphPad Inc., La Jolla, CA). Statistical tests are identified for each method. Significance was considered as p<.05 Plotting was performed using R version 4.0.4 with the following packages: ggplot2, dplyr, reshape2, ggsignif, ggpubr and plotrix.

## Author Contributions

E.M. Greeson and C.S. Madsen contributed equally to experimental design, performing experiments, data analysis and manuscript writing. A.V. Makela contributed to experimental design, performed *in vivo* studies and contributed to manuscript writing. C.H. Contag contributed to experimental design and data analysis, and manuscript writing. . All authors have given approval to the final version of the manuscript. ‡E.M. Greeson and C.S. Madsen contributed equally.

## ACKNOWLEDGMENT

The authors would like to acknowledge A. Withrow, C. Flegler, and S. Flegler at the MSU Center for Advanced Microscopy, the MSU Flow Cytometry Core, S. Rebolloso at the MSU Veterinary Diagnostic Laboratory, L. Kroos, M. Witte, A. Tundo, and K. Conner at MSU, M. Huebner and J. Tait at MSU Center for Statistical Training and Consulting. The graphical abstract and Figure 1 were created using Biorender.com.

