## Supporting Information for "Magnetothermal control of temperature-sensitive repressors in superparamagnetic iron nanoparticle-coated *Bacillus subtilis*"

Figures S1–S6 with additional experimental results and Tables S1–S2, containing additional experimental results and a list of oligos used (PDF).

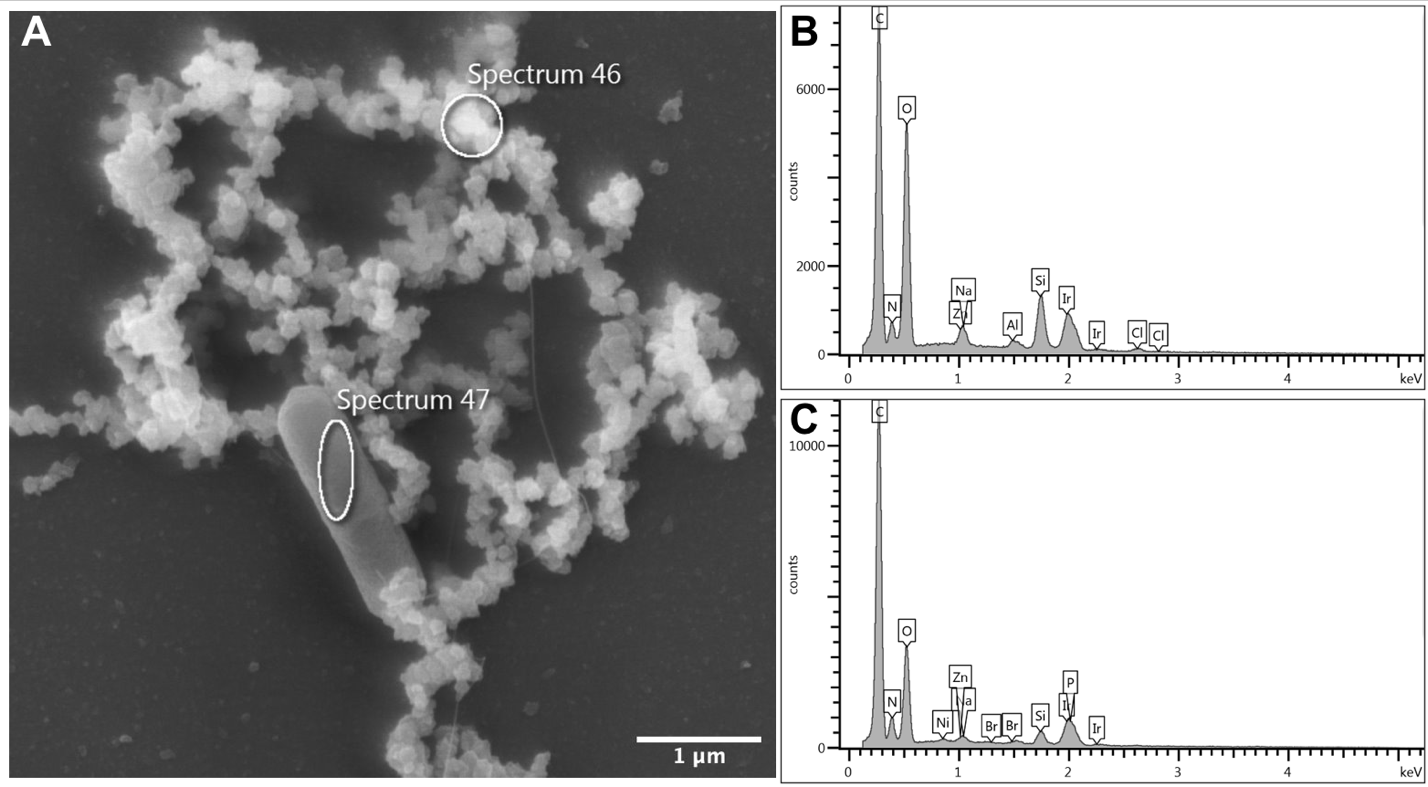

**Figure S1.** Elemental analysis of *B. subtilis* uncoated control. Sample analyzed with SEM-EDS in two regions (A) showing no iron (Fe) signal from the extracellular material (B) or from the bacterium (C). Magnification = 12,000X; scale bar = 1 µm.

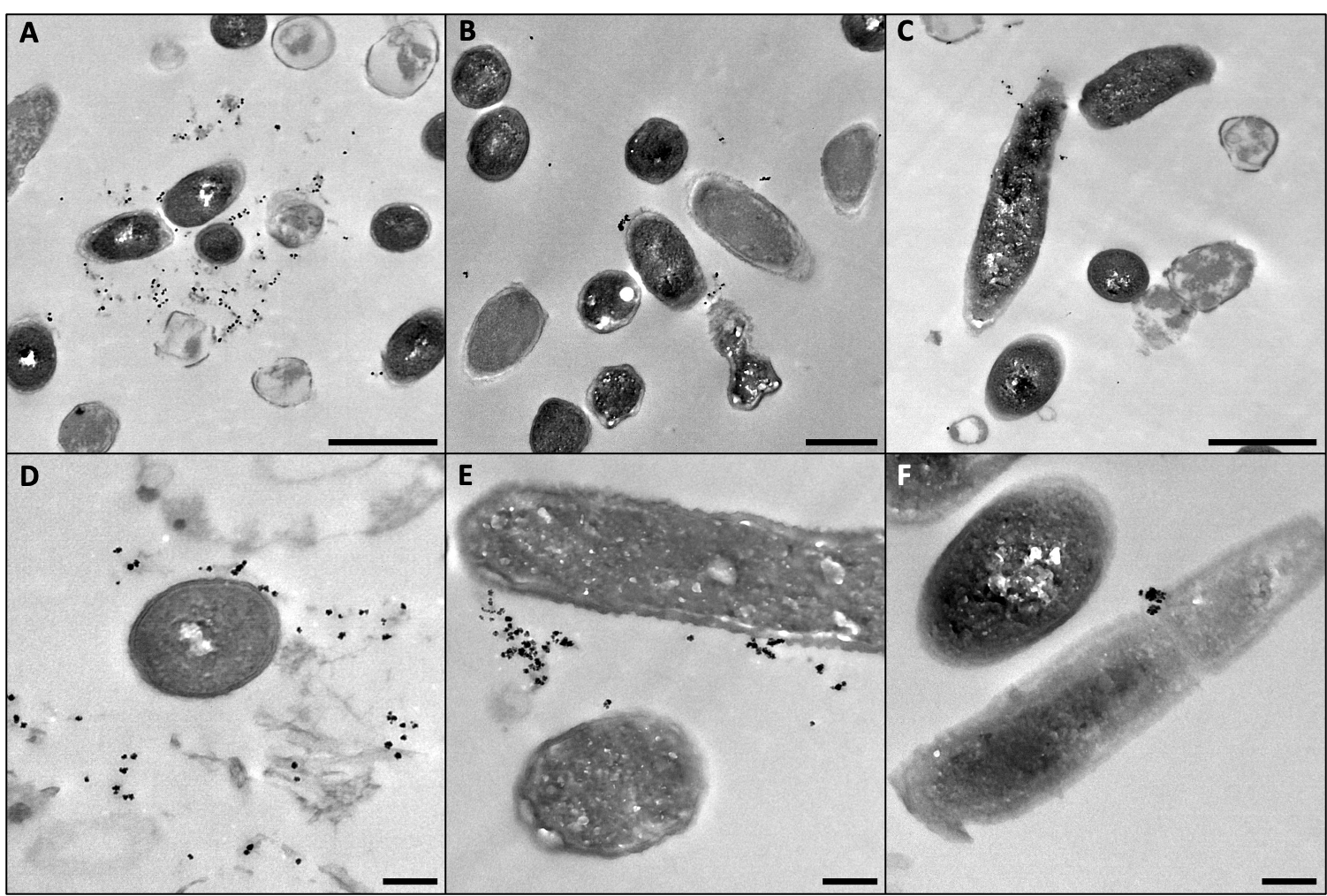

**Figure S2.** Transmission electron microscopy of *B. subtilis* coated with three SPIONs. Embedded and cross-section of each *B. subtilis*-SPION coating variation: plain-dextran (A, D), carboxyl (B, E), and amine (C, F) showing nanoparticles located outside of the bacterial cells. Mag. = 8,000X; scale bars = 1 µm (A-C). Mag. = 20,000X; scale bars = 200 nm (D-F).

**Table S1.** Quantification of iron via ICP-MS. Each of the three SPION variations and an untreated control sample were analyzed via ICP-MS. Values for the three experimental samples were adjusted by subtracting the background iron present from the untreated sample. n = 1 for all samples.

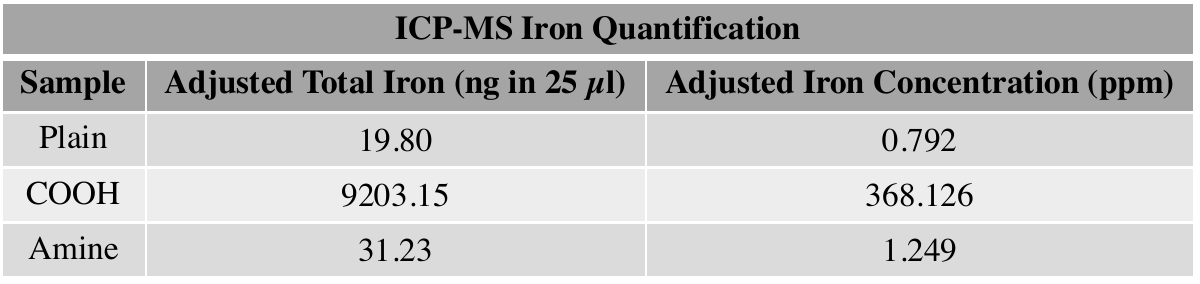

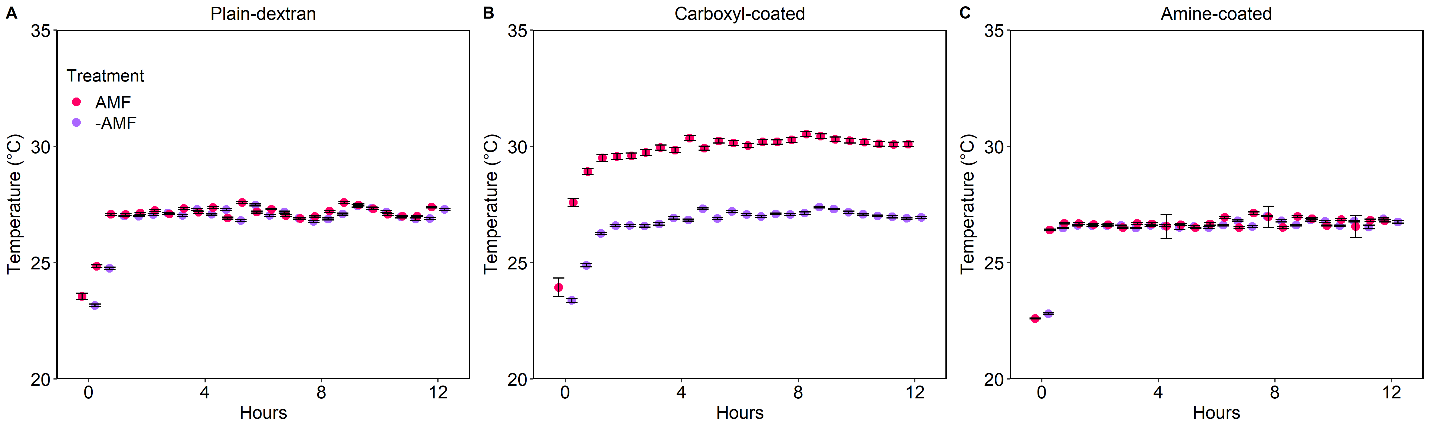

**Figure S3.** Magnetic hyperthermia impact on culture medium temperature with three Synomag-D variations. Points indicate temperature every 0.5 h over 12 h with 60 temperature reads taken every 1 m cycle considered as technical replicates. Error bars are mean ± standard deviation.

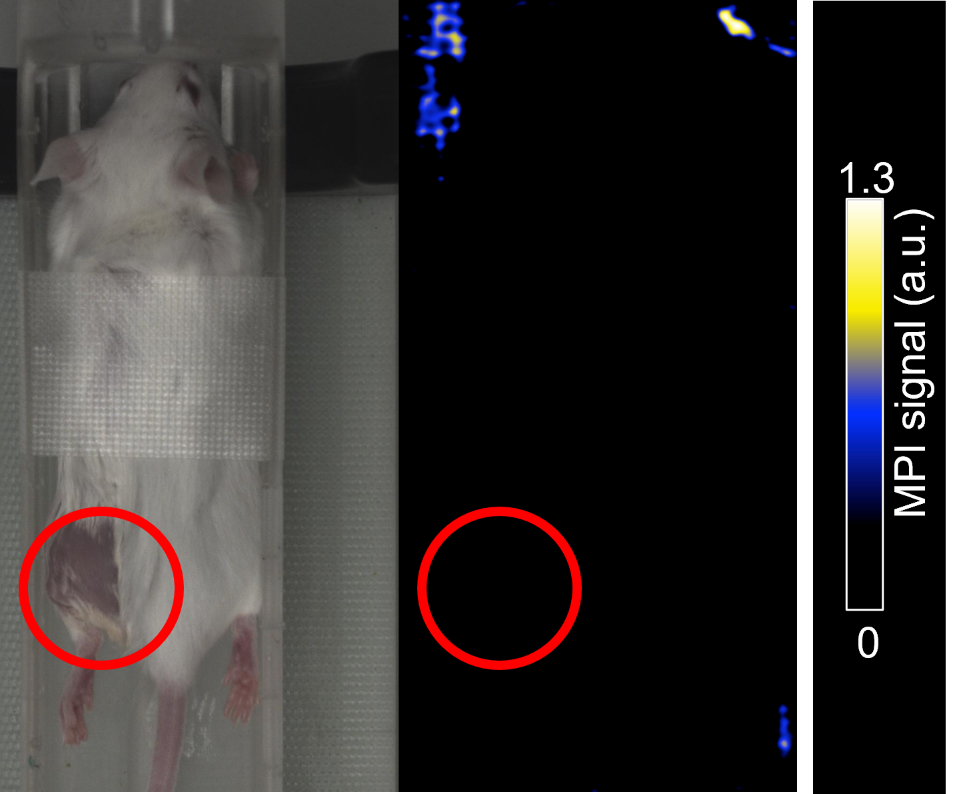

**Figure S4.** MPI of mouse thigh shows no signal from plain-dextran SPION. After injection of plain-dextran coated *B. subtilis* the -AMF control mouse was imaged via MPI and no discernible signal above background was found. Because of the injections of SPION-coated bacteria being below the limit of detection no additional mice were imaged (n=1). a.u. = arbitrary units.

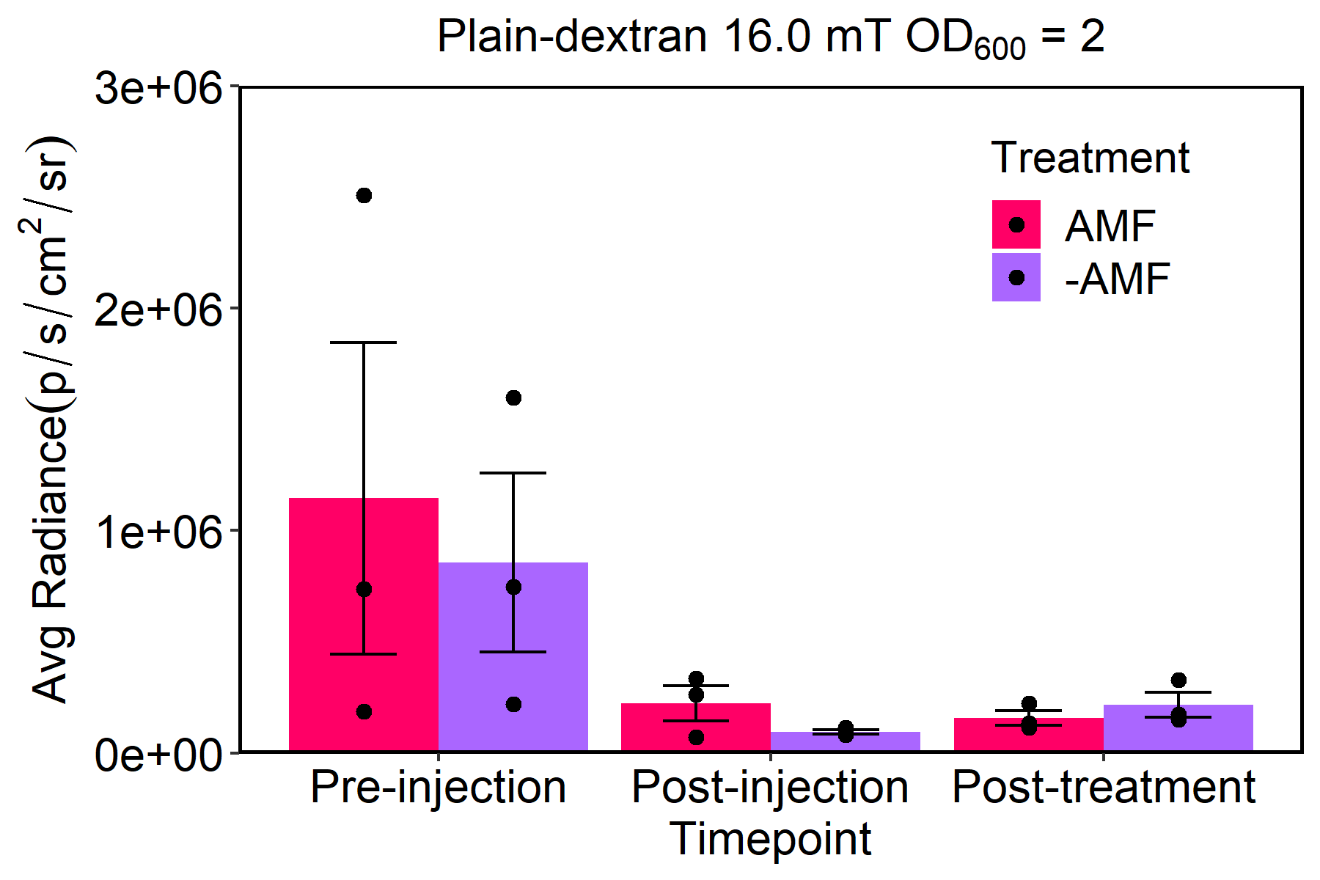

**Figure S5.** Magnetic hyperthermia of intramuscular injections did not significantly change bioluminescent signal (Avg. Radiance) when using the plain-dextran SPION. *B. subtilis* +TlpA39R was compared in and outside the AMF after injection and after 1 h of magnetic hyperthermia. Error bars are mean ± standard error mean; n = 3; two-way repeated measures ANOVA with Tukey’s post hoc showed no significance for any comparisons.

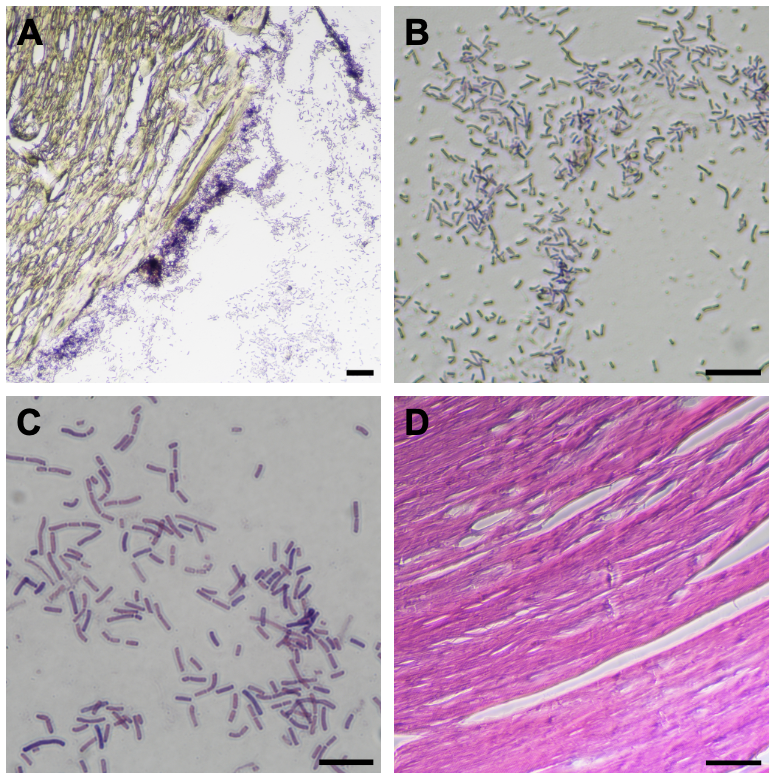

**Figure S6.** Modified Gram stain and hematoxylin and eosin staining of tissue sections. Mouse thigh muscles were sectioned and subsequently stained with a modified Gram stain method. Samples were imaged at various magnifications: 100X (A), 400X (B), and 1000X (C) with scale bars of 50 µm, 25 µm, and 10 µm, respectively. Muscle tissue is yellow from the alcoholic saffron counterstain and the purple rods are Gram positive from crystal violet which supports identification as *B. subtilis*. Hematoxylin and eosin staining was performed on mouse thigh controls with no samples injected. Imaging showed pink, eosin-stained longitudinal view of quadriceps muscle fibers; Magnification = 400X; scale bar = 25 µm (D).

**Table S2.** List of Oligos. These primers were used for assembly of the constructs described in this study as well as to measure gene expression via RT-qPCR. Refer to the methods section for specific details.

| **Oligo List** | |
| --- | --- |
| **Primer** | **Sequence (5’-3’)** |
| pDR111_linear_hyper_F | ctgcataaaaaacgcccggc |
| pDR111_linear_hyper_R | taagcttagtcgacagctagccg |
| TlpA39_promoter_F | gccgggcgttttttatgcagctcgatcccgcgaaatttta |
| TlpA39_promoter_R | cggctagctgtcgactaagcttatggatcctggctgtggt |
| pDR111_linear_LacI_F | cggctagctgtcgactaagctta |
| pDR111_linear_LacI_ R | tgagttaggatcctgagcgcc |
| TlpA39_regulator_F | taagcttagtcgacagctagccgatgagctcgacaccatc |
| TlpA39_regulator_R | ggcgctcaggatcctaactcatcctcctttcagcaaaaaac |
| LuxAE_TlpA39_promoter_overhang_F | ccgaattagcttgcatgcggttaactatcaaacgcttcggtta |
| LuxAE_TlpA39_promoter_overhang_R | gatccataagcttagtcgacagatgaagcaagaggaggactctctatg |
| LuxAE_TlpA39_system_overhang_F | cagtaagcttagtcgacagatgaagcaagaggaggactctctatg |
| LuxAE_TlpA39_system_overhang_R | cgatggtgtcgagctcatcggggttaactatcaaacgcttcggtta |
| TlpA39R_qPCR_F | ggccttcacttcttcagcca |
| TlpA39R_qPCR_R | acggaatatcaccgggttcg |
| luxB_qPCR_F | taatggtgttgtcggcgct |
| luxB_qPCR_R | aataagcaagcttcctccgct |
| 16S_qPCR_F | ggtaatggcctaccaaggca |
| 16S_qPCR_R | tgctccgtcagactttcgtc |
